# Adaptive Differentiation in the General-Purpose Genotype Invasive Plant *Erythranthe guttata*

**DOI:** 10.1101/2024.04.30.591966

**Authors:** Aaron O Millar, Hazel M Chapman

## Abstract

1. Highly plastic general-purpose genotypes are a frequent occurrence among invasive plants. Yet, it remains uncertain to what extent genetic adaptation can co-occur with such elevated levels of plasticity. Understanding the interplay between these two evolutionary strategies is essential to better predict invasive success and future climate change responses.
2. We investigated the potential for local adaptation along an altitudinal gradient in introduced New Zealand populations of the highly plastic invasive herb, *Erythranthe guttata*. We asked a) whether there were phenotypic differences between upland and lowland *E. guttata* populations along our gradient; b) whether any differences were consistent with known adaptive patterns; and c) whether any adaptive patterns exist alongside high plasticity to elevation.
3. Samples from 38 *E. guttata* populations from upland and lowland Canterbury were grown from cuttings in a lowland and an upland common garden, where we measured a broad range of growth and reproductive traits.
4. We found significant adaptive differentiation between upland and lowland populations over almost all measured traits. Upland populations had earlier and more intense flowering compared with lowland populations. Lowland plants were taller and had larger leaves with higher photosynthetic rates than upland plants. These differences occurred alongside high levels of unspecialised plasticity to the growing environment.
5. Synthesis: We found that over a period of less than 150 years the environment along an altitude gradient of 120km has selected for distinct lowland and upland phenotypes of *E. guttata*. These changes reflect common selective pressure associated with altitude gradients, increasing reproductive success at higher altitudes and increased competitive ability at lower altitudes. This rapid local adaptation occurred alongside high plasticity within the growing environment, showing that highly plastic invasive species still retain the capacity to genetically adapt to novel environments.

## 1: Introduction

Many globally widespread invasive species have highly plastic ‘general-purpose genotypes’ (Baker, 1965; Bolpagni, 2021; Cornille *et al*., 2022) which respond to environmental gradients and stressors primarily through phenotypic plasticity (Geng *et al*., 2007; Matesanz *et al*., 2012; Bolpagni, 2021). The term general-purpose genotype also implies a lack of genetic adaptation, as such species usually show very little genetic differentiation or local adaptation (Geng *et al*., 2007; Ross *et al*., 2009; Riis *et al*., 2010; Matesanz *et al*., 2012, Querns *et al*., 2022). However, genetic differentiation may still be possible in these species (Williamson *et al*. 2023) and identifying to what extent it occurs will have major implications for understanding current and future invasive success (Bolpagni, 2021).

Local adaptation is common and often rapid in invasive species (Odour *et al*., 2016). For this reason, invasive species have been used as a model for understanding rapid adaptation (Clements & Jones, 2021) and identifying the early emergence of adaptive divergence (Prentis *et al*., 2008; Lucek *et al*., 2013). Local adaptation has the most evolutionary impact when acting on suites of traits at once (Lucek *et al*., 2013, Felmy *et al*., 2022). Because selection on suites of traits represent success strategies, they are important for identifying selective gradients acting on invasive species (Radford & Cousens, 2000). In invasive species these adaptations often occur over similar selective gradients and in parallel traits to the native range (Lucek *et al*., 2013; Latimer *et al*., 2019; Hernández *et al*., 2019; Van Boheemen *et al*., 2019; McGoey *et al*., 2020).

There are, however, many examples of high plasticity in invasive plants preventing adaptive divergence (Van Kleunen *et al*., 2011; Matesanz *et al*., 2012; Li *et al*., 2017; Matesanz *et al*., 2020; Van Wallendeal *et al*., 2021; Villelas *et al*., 2021; Cornille *et al*., 2022; Querns *et al*., 2022). Adaptive differentiation requires that selection is sufficiently strong to outweigh the homogenising effect of gene flow (Schmidt & Garroway, 2018). Phenotypic plasticity allows species to produce phenotypes to match different environments without genetic change (Fusco and Minelli, 2010). Having well-matched phenotypes reduces the selective gradient and so can prevent genetic adaptation (Villellas *et al*., 2021). Furthermore, in highly plastic invasive species local adaptation often acts on plasticity levels themselves, rather than trait means (Lande, 2015; Coluatti *et al*., 2017; Kelly, 2019, Matesanz *et al*., 2020; Bufford & Hulme, 2021a). Highly plastic introduced species therefore have major barriers to mounting adaptive phenotypic responses. However, this is not inevitable; plasticity may facilitate selection by allowing survival in mismatched environments while slower selection processes occur (Matesanz *et al*., 2010; Nicotra *et al*., 2010, Levis & Pfennig, 2016). Plasticity can also synergistically aid adaptive divergence, when genetic adaptation and plasticity act on complementary traits (Lucek *et al*., 2014).

It is important to understand the capacity for general-purpose genotypes to show adaptive genetic change alongside high plasticity levels. Local adaptation is an important potential driver of invasive success (Bock *et al*., 2018). Rapid evolution is also anticipated to play a major role in how invasive plants will respond to climate change (Richardson *et al*., 2014; DeMarche *et al*., 2020; Clements & Jones, 2021), while plastic responses alone may not be enough (Franks *et al*., 2014). Identifying to what extent local adaptation is possible for general-purpose genotype invaders is important for predicting their future invasiveness. To understand this requires investigation of both the plastic and genetic components of adaptive differentiation in these species (Felmy *et al*., 2022). Adaptative and plastic responses can differ among traits (DeMarche *et al*., 2016; Villelas *et al*., 2021), so a wide range of traits must be assessed at once, over an ecologically relevant gradient (Forsman, 2015; Popovic & Lowry, 2020; Schneider, 2022).

An ideal system for studying the potential for local adaptation in general-purpose genotype invaders is the yellow monkeyflower *Erythranthe guttata* (also referred to as *Mimulus guttatus*) invading the Canterbury region of New Zealand. *E. guttata* is an evolutionary model species (Braindvain *et al*., 2014) from North America. As an invasive species it has a general-purpose genotype, with high levels of plasticity and little evidence of adaptive patterns (Holeski, 2007; Querns *et al*., 2022; Williamson *et al*., 2023). However, in its native range *E. guttata* shows large amounts of local adaptation (Friedman *et al*., 2015; Kooyers *et al*., 2019), especially based on altitude (DeMarche *et al*., 2016; Popovic & Lowry, 2020). Local adaptation has been identified in a few traits and genes in its invasive United Kingdom range (Puzey & Vallejo-Marín, 2014; Simón-Porcar *et al*., 2021), but this adaptation has not occurred in consistent suites of traits over environmental gradients (Simón-Porcar *et al*., 2021).

*Erythranthe guttata* has been naturalised in New Zealand since 1878 (Vallejo-Marín *et al*., 2021). The Canterbury region consists of lowland plains where *E. guttata* is very well established (below 100 metres above sea level), and a mountainous core where it is actively expanding its range and increasing in abundance (above 450 masl) (Williamson *et al*., 2023). The total altitudinal gradient extends from 0-900 masl over distances of ∼120km. This altitudinal gradient is a potential site of adaptive differentiation in *E. guttata* populations (DeMarche *et al*., 2016; Kooyers *et al*., 2019; Popovic & Lowry, 2020). Da Re *et al*. (2020) suggested that genetic adaptation to this gradient may be required for *E. guttata*’s successful spread into upland Canterbury. However, research into latitudinal variation in New Zealand populations of *E. guttata* found minimal genetic variation, with success explained mostly by high plasticity (Williamson *et al*., 2023). This potential for local adaptation in a species well-known for its general-purpose genotypes makes New Zealand *E. guttata* a most suitable study plant to understand how these two evolutionary dynamics may interact.

A strong local adaptive response would point to adaptive differences in suites of traits associated with known selective drivers (Lucek *et al*., 2014). Altitudinal gradients can be steep, with more challenging abiotic conditions and greater seasonality at high altitudes (Rubin & Friedman, 2018), and higher levels of competition at lower altitudes (Choler *et al*., 2001). These diverging pressures select for reproductive success at high altitudes and greater competitive ability at lower altitudes (Bodbyl Roels & Kelly, 2011; Fenster & Ritland, 1994). This is commonly observed as diverging reproductive and growth traits (Choler *et al*., 2001; van Kleunen & Fischer, 2008). Yet another trait that may be evolving in Canterbury *E. guttata* is self-pollination at higher altitudes, which has been observed occurring based on latitude in invasive British populations of *E. guttata* (Puzey & Vallejo-Marín, 2014). It has also been linked to pollinator shortages in upland environments in *E. lutea* (Carvallo & Medel, 2009). These adaptations occur as suites of changes called selfing syndromes, including decreased flower size, reduced herkogamy (anther-stigma separation), and reduced vegetative mass (Bodbyl Roels & Kelly, 2011; Razanajatovo *et al*., 2011).

This study asked a) whether there are phenotypic differences between upland and lowland Canterbury *E. guttata* populations; b) whether any differences are consistent with known adaptive responses to altitudinal variation and c) whether any adaptive patterns exist alongside high plasticity to elevation.

## 2: Materials and Methods

### 2.1: Sampling Populations

We sampled 278 individual plants from 38 yellow monkeyflower populations from 5 upland and 5 lowland Canterbury river systems in October and November 2021 (Supplementary Table S1). These locations were chosen to reduce the possibility of genetic mixing through downstream spread (Truscott *et al*., 2007). Populations were sampled from a wide variety of habitats including streams, roadside drains, and farm culverts. 2-10 plants per population were sampled depending on population size, with each waterway system sampled with equal frequency.

At the University of Canterbury, we made four 10-40mm long cuttings from each plant from a mix of internodes and young shoot tips. Short cutting size minimises maternal effects (Williamson *et al*., 2023). We placed the four cuttings from each plant together in small pots filled with fertilised potting soil and placed them under a mister where they were kept constantly warm and wet. We thinned out duplicate cuttings as the plants grew, until only one individual cutting per plant remained. Cuttings were transplanted into common gardens when they reached approximately 10cm in height.

### 2.2 : Common Gardens

We used two common gardens to identify adaptive (Popovic & Lowry, 2020) and plastic (Matesanz *et al*., 2010) variation, a lowland garden at the University of Canterbury (5 masl) and an upland garden at the Cass field station (583 masl). These locations were chosen to reflect the climate conditions in lowland and upland Canterbury respectively. The lowland garden housed clones of all 278 plants and was established in November-December 2021. The upland garden held a representative subset of 120 plants produced from a second round of cuttings taken from the lowland garden. These received two weeks of outdoor acclimation prior to planting out to minimise damage from temperature shock. Due to weather and facility complications the upland garden was established in early October 2022. In both gardens plants were housed in 7-litre pots and placed in a randomised block design. They were watered amply once per day by an automatic watering system and fertilised prior to the research period.

### 2.3 : Data Collection

Data collection occurred from the end of October 2022 to April 2023. We measured 13 vegetative and reproductive traits already known to show adaptive and plastic variance in *E. guttata* (Galloway, 1995; DeMarche *et al*., 2016; Simón-Porcar *et al*., 2021; Villellas *et al*., 2021; Williamson *et al*., 2023). For vegetative traits, above-ground biomass was measured as dry weight. All above-ground growth was harvested in April 2023, at the end of the regrowth period after summer flowering and then dried for 5 days at 80°C before being weighed. We did not measure below-ground biomass because of root binding in pots. Mean leaf length and width were calculated from the three largest leaves on each plant. Plant height was measured during the fifth week of flowering. Photosynthetic rates were measured for three young leaves on each plant using an Infrared Gas Analyser (LI-COR Biosciences, model LI-6400). Measurements were taken using external light on fine summer days over 20°C between the hours of 10am and 4pm. For reproductive traits, flower number was counted from dried seedheads after flowering. Seed mass produced per flower, average seed mass, and seed number per flower were measured for each plant. Three seed pods per plant were collected from separate stems just before seed release during peak seeding and dried at room temperature in paper envelopes for at least four weeks. The seeds were weighed on scales to calculate seed mass per flower. Around 50 seeds were also separated out, counted, and weighed on a microbalance. This measure was conducted twice for each plant and used to calculate average seed mass and seed number. Flower corolla width, a proxy for flower size in *E. guttata* (Kleunen & Ritland, 2004), and herkogamy were measured using digital callipers. First flower date and total bloom period were documented. *E. guttata* flowering involves a long tail of individual flowers produced well after the main floral display has ended, so the end of flowering was defined as the first day when all major upright flowering stems had no open flowers or live buds. The later establishment date of the upland garden meant accurate phenological data could only be collected in the lowland garden. Outside of the growth and reproductive measures, the level of anthocyanin colouration in stems and leaves was measured qualitatively, as it is known to vary based on plant stress levels (Twyford *et al*., 2018) and genetics (Twyford *et al*., 2020). A late frost also damaged young plants in the upland garden, allowing a qualitative assessment of the level of cold damage suffered to be measured.

### 2.4 : Insect Visitor Observations

Flower visitor observations were used to assess whether *E. guttata* was experiencing pollinator limitation in upland, compared to lowland Canterbury. Though not perfectly accurate (Arroyo *et al*., 1985), flower visitor rates are a reliable proxy for pollinator activity (Minachilis *et al*., 2021; Totland, 2001). Observations were conducted over multiple sites in five upland and five lowland Canterbury waterway systems in January 2021. Bees are the primary pollinators of *E. guttata* in its native range (Bodbyl Roels & Kelly, 2011), so observations were conducted in condition suitable for bees: temperatures above 12°C, low wind speeds, and non-overcast skies (Ullman *et al*., 2008). Multiple patches of 50-70 flowers were observed in 15 minutes blocks throughout the day. Insects were counted as pollinators if they entered the corolla tube.

### 2.5 : Data Analysis

The effect of origin and garden location on phenotype was analysed using linear mixed effects models. Population and the individual plants nested within them were used as random effects. Hypothesis tests were conducted through analysis of deviance using Type II Wald chisquare tests. Coefficients of determination were calculated using the MuMIn package when significance was identified. A significant interaction term between origin and garden location was interpreted as a difference in phenotypic plasticity levels between upland and lowland *E. guttata* populations. Where assumptions of the tests were not met, appropriate data transformations or non-parametric equivalents were used. Flower number was analysed with bloom period controlled for. Seed traits and herkogamy were analysed with corolla width controlled for as a proxy for flower size (Kleunen & Ritland 2004). Photosynthetic rates were analysed with the time of day the measurement was taken controlled for to account for differences in temperature and light intensity. Effect sizes were also calculated using Hedge’s G, which serve as another measure of assessing significant differences and plasticity levels (Stotz *et al*., 2021). Insect visitor frequency was compared between upland and lowland areas using a poisson distribution in a generalised linear model. Mixed effects models were also used to check for possible trade-offs between pairs of traits. All statistical analyses were conducted using R statistical software version 4.2.0 (R Core Team, 2022).

## 3: Results

### 3.1: Phenotypic Differences Between Lowland and Upland Erythranthe guttata

We identified significant phenotypic differences in yellow monkeyflower plants from upland and lowland areas in most traits (Fig. 1, Table 1). Upland *E. guttata* populations flowered earlier and had increased flower numbers compared to lowland populations, producing 25% more flowers on average in the lowland garden, flowering 15 days earlier, and for 7 days longer (Supplementary Table S2). Upland populations also experienced less cold damage than lowland populations. Upland *E. guttata* populations also had significantly less leaf and stem anthocyanin colouration than lowland populations in both gardens. In contrast, lowland populations had higher trait means than upland populations in most growth traits (Fig. 1). Photosynthetic rate was 15% higher on average in lowland *E. guttata* populations than upland, a significant difference that was seen in both gardens (Supplementary Table S2). Lowland *E. guttata* populations grew 24% taller on average in the lowland garden, though there was no difference in the upland garden (Supplementary Table S2). Lowland populations had leaves that were 16-17% wider than upland populations in both gardens and 8% longer in the lowland garden. Origin explained more than 20% of observed variation in three traits: cold damage experienced, first flower date, and leaf anthocyanin levels (Table 1). Significant effects of origin in other traits were weak; in most cases origin explained under 10% of the measured variation, and often under 5% (Table 1). Dry weight, flower width, herkogamy, and all seed traits showed no differences between origin groups in either garden.

**Figure 1:**
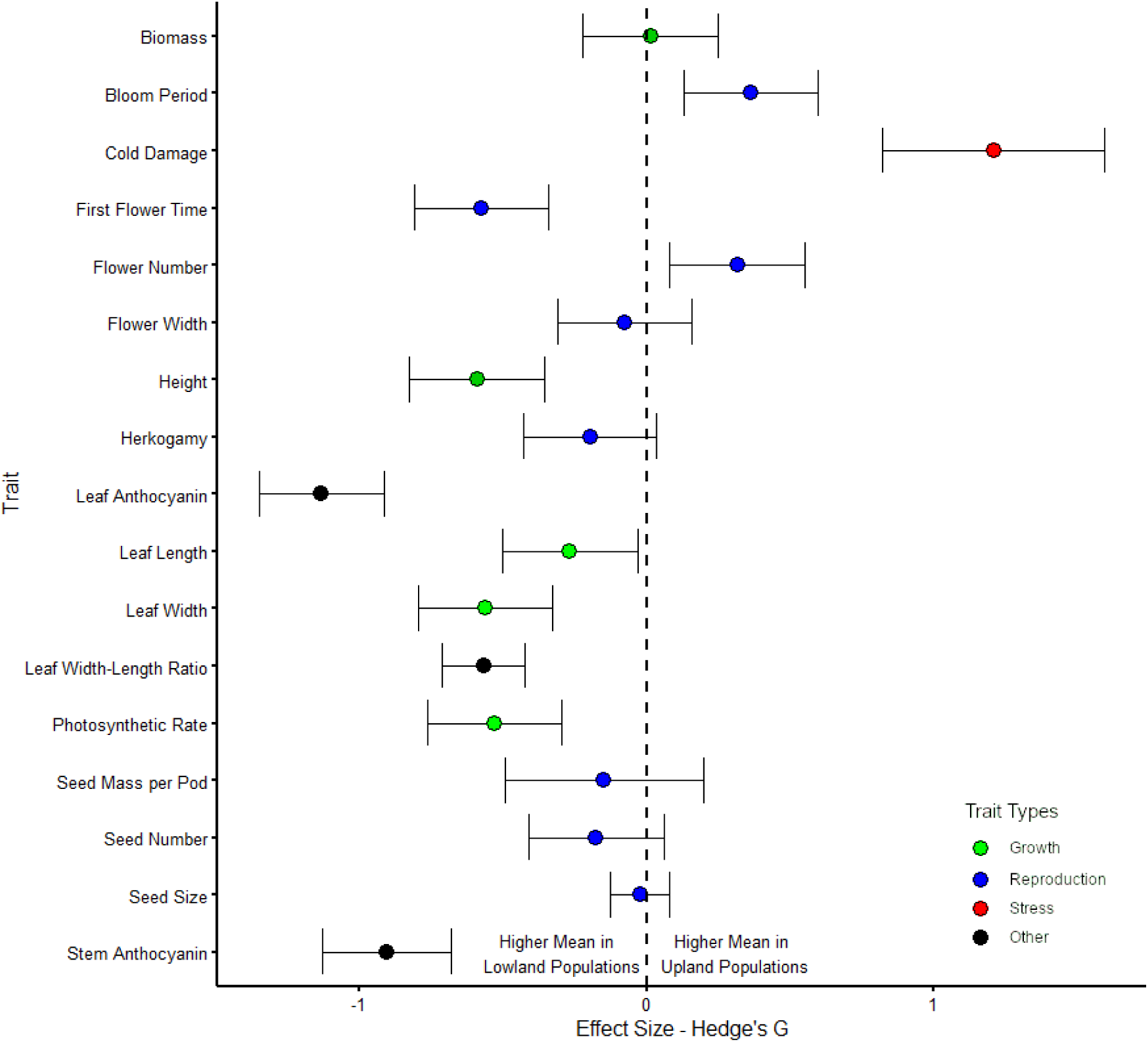
*Hedges G effect sizes and 95% confidence intervals for the effect of altitudinal origin on trait phenotype in Canterbury* Erythranthe guttata *populations with traits coloured by type*.

**Table 1:** Analysis of deviance results from type II Wald chi-square tests on mixed effects models comparing the effect of altitudinal origin and growing location on phenotype in Canterbury Erythranthe guttata populations. Significant items are shown in bold.

| Trait | Controls | Origin<br>Chisq | Origin<br>P Value | R<sup>2</sup><br>Value | Garden<br>Chisq | Garden<br>P Value | R<sup>2</sup><br>Value | Garden<br>x Origin<br>Chisq | Garden x<br>Origin<br>P Value | R<sup>2</sup><br>Value |
| --- | --- | --- | --- | --- | --- | --- | --- | --- | --- | --- |
| Bloom Period |  | **7.9936** | **0.004694**<br>** | **0.04** |  |  |  |  |  |  |
| Cold Damage |  | **19.77** | **8.734e-06**<br>*** | **0.24** |  |  |  |  |  |  |
| Corolla Width |  | 0.1745 | 0.6761 |  | 0.0592 | 0.8077 |  | 0.8382 | 0.35992 |  |
| Dry Weight |  | 0.0846 | 0.77119 |  | 0.9776 | 0.3228 |  | 1.4774 | 0.22419 |  |
| First Flower<br>(Days from<br>Start 2022) |  | **18.1437** | **2.048e-05**<br>*** | **0.29** |  |  |  |  |  |  |
| Flower<br>Number | Bloom<br>Period | **4.5586** | **0.032754**<br>* | **0.19** | **151.2391** | **< 2.2e-16**<br>*** | **0.28** | **5.8575** | **0.015511**<br>* | **0.22** |
| Height |  | **14.2711** | **0.0001583**<br>*** | **0.1** | **203.7013** | **< 2.2e-16**<br>*** | **0.25** | **20.7643** | **5.194e-06** *** | **0.33** |
| Herkogamy | Corolla<br>Width | 1.1365 | 0.286389 |  | **10.5753** | **0.001146**<br>** | **0.02** | 0.0439 | 0.834121 |  |
| Leaf<br>Anthocyanin<br>Score |  | **33.8689** | **5.895e-09** *** | **0.26** | 3.6726 | 0.055315 |  | 1.0991 | 0.294474 |  |
| Leaf Length |  | **5.8599** | **0.01549 \*** | **0.02** | **520.6254** | **< 2.2e-16**<br>*** | **0.47** | 0.5548 | 0.46044 |  |
| Leaf Width |  | **27.0105** | **2.024e-07 \*\*\*** | **0.07** | **345.8587** | **< 2.2e-16**<br>*** | **0.38** | 2.2116 | 0.136979 |  |
| Leaf Width-Length Ratio |  | **13.4178** | **0.0002492 \*\*\*** | **0.08** | **44.6797** | **2.321e-11**<br>*** | **0.07** | 0.0005 | 0.981974 |  |
| Photosynthetic<br>Rate | Time of<br>Day | **15.1887** | **9.728e-05 \*\*\*** | **0.06** | **14.8971** | **0.0001135 \*\*\*** | **0.01** | 0.5052 | 0.477212 |  |
| Seed Mass per<br>Pod | Corolla<br>Width | 0.7902 | 0.37405 |  | 1.6198 | 0.20311 |  | 2.8457 | 0.09162 . |  |
| Seed Number | Corolla<br>Width | 0.7275 | 0.3937 |  | 1.3172 | 0.251088 |  | 0.8322 | 0.3616 |  |
| Seed Size |  | 0.0027 | 0.9582 |  | 0.2362 | 0.627 |  | 0.5788 | 0.4468 |  |
| Stem<br>Anthocyanin<br>Score |  | **20.7938** | **5.115e-06 \*\*\*** | **0.18** | **15.0556** | **0.0001044 \*\*\*** | **0.02** | 2.5859 | 0.107816 |  |

### 3.2 Plastic Reponses to Growing Environment

Many traits showed plastic differences between plants in the lowland garden and the clonal subset in the upland garden (Fig. 2). In growth traits, *E. guttata* plants growing in the lowland garden were on average 40% taller and had 65% longer and 52% wider leaves than *E. guttata* plants growing in the upland garden (Supplementary Table S2). Notably however, dry weight was not significantly different between gardens. Reproductively, *E. guttata* growing in a lowland environment produced 109% more flowers than *E. guttata* growing in the lowland garden. In these significantly different traits growing environment explained more than 25% of observed variation (Table 1). The reproductive traits of corolla width, reproductive output per flower, average seed mass, and seed number per flower all did not vary between growing environments. Photosynthetic rate was very slightly higher in the upland garden, though in this case the difference is weak and almost certainly due to weather conditions during measurement.

**Figure 2:**
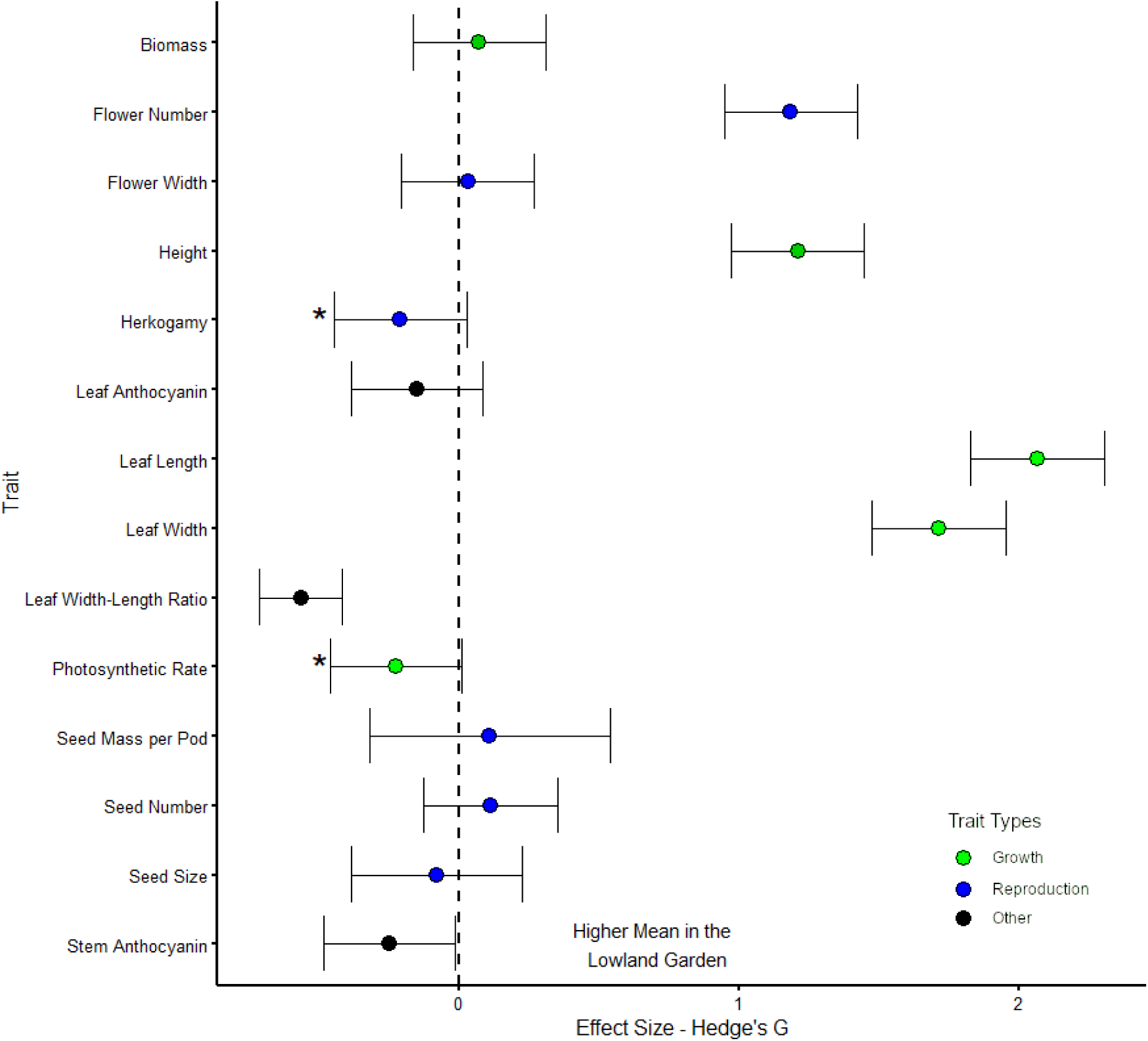
*Hedges G effect sizes and 95% confidence intervals for the effect of garden location on trait phenotype in Canterbury* Erythranthe guttata *populations with traits coloured by type. Asterisks indicate traits where garden was shown to have a significant effect on phenotypic mean in random effects models which was not reflected in the Hedge’s G calculations*.

### 3.3 Undifferentiated Plastic Responses Between Origin Groups

Upland and lowland origin *E. guttata* populations only showed differences in their plastic responses to growing location in flower number and height (Table 1). Upland origin *E. guttata* populations had strong plastic responses for flower number, with the two groups showing no difference in flower number in the upland garden, but in the lowland garden upland *E. guttata* plants produced on average 25% more flowers than lowland *E. guttata* plants (Fig. 3, Supplementary Table S2). Conversely, lowland *E. guttata* plants had greater height plasticity. Upland and lowland plants showed no difference in the upland garden, but in the lowland garden lowland origin *E. guttata* grew 24% taller on average.

**Figure 3:**
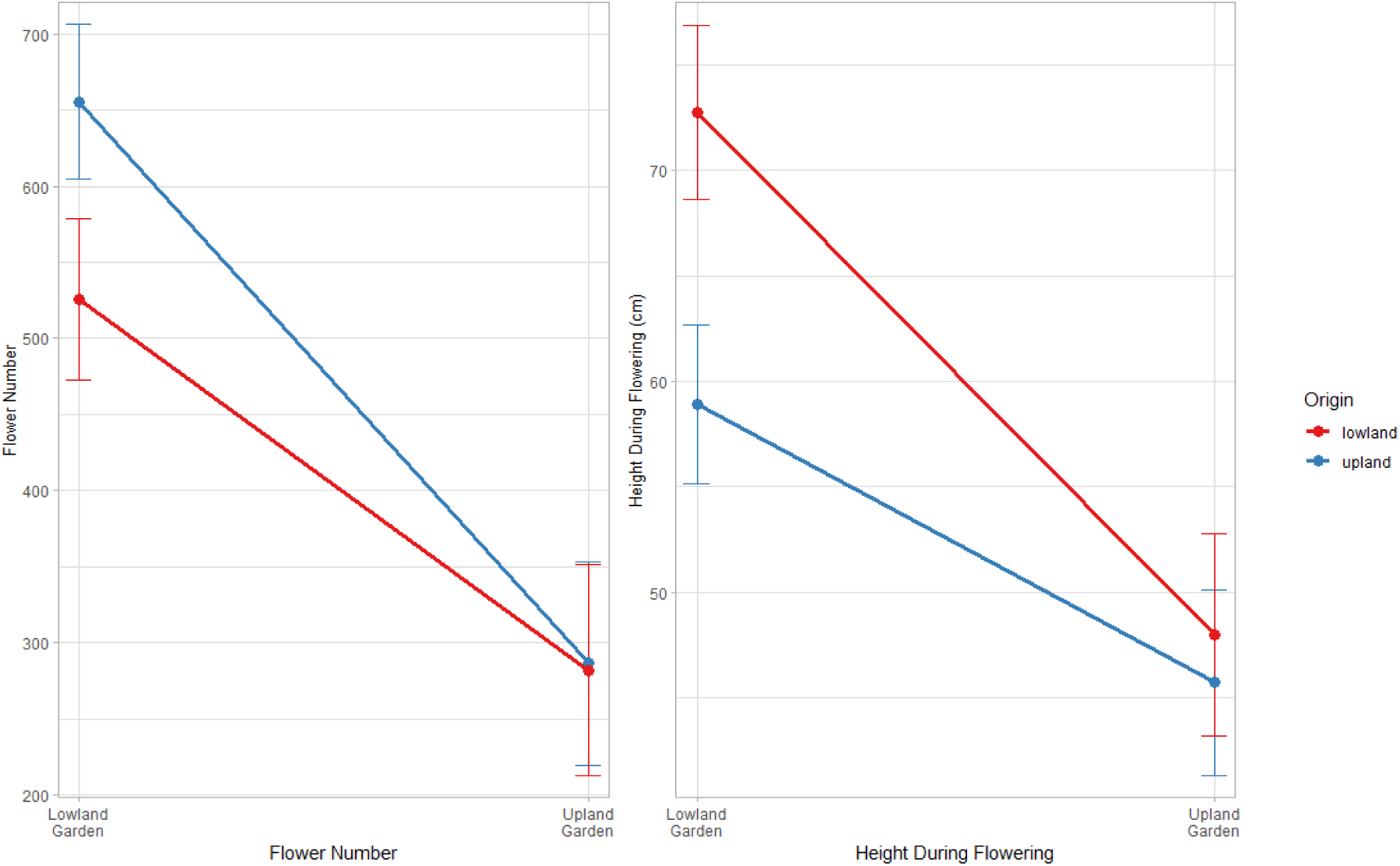
*Reaction norm graphs showing phenotypic mean and 95% confidence interval for* Erythranthe guttata *of lowland and upland origin growing in lowland and upland gardens, for traits where a significant interaction between origin and growing location on phenotype was observed in mixed effect models*.

### 3.4 Flower Visitation and Self-Pollination Related Traits

There was no significant difference in flower visitation to *E. guttata* between upland and lowland areas once accounting for outliers (Deviance_1, 351_: 0.00047603, P: 0.98; Supplementary Fig. S1). Insect visitor species composition did change between sites, however. Native *Leioproctus fulvescens* and *Melangyna novaezalandiae* were more common visitors in upland areas than lowland areas and were the majority of visitors to *E. guttata* during observations in the Lake Ohau and Lake Pukaki areas. Elsewhere introduced *Apis mellifera* and *Bombus* species were the main pollinators.

The traits associated with selfing syndromes were also not observed among Canterbury *E. guttata*. Upland and lowland populations did not differ in flower width or herkogamy, and in the upland garden herkogamy increased marginally rather than decreasing, while flower size did not vary significantly (Table 1).

## 4: Discussion

### 4.1 Altitudinal Differentiation

This experiment identified clear differences between upland and lowland Canterbury *E. guttata* populations, consistent with genetic differentiation. These differences occurred in suites of traits that correspond to known evolutionary responses to altitudinal gradients (Choler *et al*., 2001; van Kleunen & Fischer, 2008). The upland *E. guttata* phenotype is consistent with selection for reproductive success, with earlier flowering and greater reproductive output in upland populations compared to lowland populations (Johansson *et al*., 2013). Higher altitudes select for reproductive success due to more challenging abiotic conditions and greater seasonality (Johansson *et al*., 2013; Rubin & Friedman, 2018; van Kleunen & Fischer, 2008). Selection for increased reproductive success in more challenging abiotic conditions and higher altitudes has previously been observed in native range *E. guttata* populations (Galloway, 1995; DeMarche *et al*., 2020). Upland *E. guttata* populations also received less frost damage than lowland populations, a common alpine adaptation (Malyshev *et al*., 2014). Lowland *E. guttata* phenotypes favoured traits associated with more aggressive growth. High growth rates can provide competitive advantages (Fischer *et al*., 2008; Kooyers *et al*., 2017) and so are selected for at the lower end of altitude gradients due to increased competition (Choler *et al*., 2001). Competition is a major selective driver in *E. guttata* (Truscott *et al*., 2008), and this finding is consistent with faster growth rates being important to lowland *E. guttata* populations in the native range (Kooyers *et al*., 2019). The lack of biomass variation is contrary to previous findings in *E. guttata* (Kooyers *et al*., 2019; Simón-Porcar *et al*., 2021), but may be due to dry weight measurements being made in autumn rather than during the peak growth period. While variation between origin groups often explained little overall trait variation, this may be due to high levels of variation in *E. guttata* overall (Twyford *et al*., 2020, Williamson *et al*., 2023) or differentiation being still emerging (Prentis *et al*., 2008). Our findings demonstrate that known selective responses to altitude in *E. guttata*’s native range are being replicated during the Canterbury invasion (Angert, 2006; Hall *et al*., 2010; Friedman *et al*., 2015; DeMarche *et al*., 2016; Kooyers *et al*., 2019). While earlier research into New Zealand *E. guttata* did not detect altitudinal differentiation (Williamson *et al*., 2023), this new clarity may be explained by strong local sampling overcoming chaotic local scale variation in *E. guttata* (Twyford *et al*., 2020). It is notable however, that adaptations for increased self-seeding in some populations were not observed. The lack of pollinator shortages suggest that the selective gradient required to drive these adaptations is not present in Canterbury (Van Kleunen & Fischer, 2008).

Differentiation between upland and lowland Canterbury *E. guttata* populations is more consistent with genetic change than phenotypic plasticity. Significant differences between origin groups frequently occurred in both upland and lowland gardens, suggesting a genetic basis (Savolanien *et al*., 2013). Nor did the pattern of phenotypic differentiation between origin groups match their plastic responses to the same environmental gradient. Phenotypic differences occurred between origin groups in most traits, whereas plastic responses to growing location were largely uniform between origin groups, suggesting these dynamics were not linked (Schneider *et al*. 2022). This is consistent with past findings that most trait shifts in *E. guttata* are genetic (Galloway, 1995; DeMarche *et al*., 2020; Williamson *et al*., 2023). *E. guttata* in New Zealand has a history of multiple introductions and high diversity that could facilitate genetic differentiation (Riis *et al*., 2010; Vallejo-Marín *et al*., 2021). Whether this differentiation specifically represents local adaptation is less clear. Evidence for local adaptation ideally demonstrates evidence of local genotypes having higher fitness in their respective environments (Kooyers *et al*., 2019). We instead found upland genotypes having higher fitness in both gardens, in the form of greater reproductive output. However, these counterintuitive fitness findings are not uncommon in *E. guttata* (DeMarche *et al*., 2016; Williamson *et al*., 2023). They are suggested to be due to experiments not capturing all selection factors in the environment, with local genotypes expected to be more fit once less frequent environmental stressors are accounted for (DeMarche *et al*. 2016). In this experiment, the identification of genetic differentiation over suites of traits that correspond to known adaptive patterns is strong indicative of local adaptation (Lucek *et al*., 2014). This supports the idea that genetic shifts may be required for *E. guttata*’s success in upland Canterbury (Da Re *et al*., 2020). Further research with reciprocal transplants would shed further light on this issue.

Adaptive genetic differentiation in Canterbury *E. guttata* fits into a broader pattern of parallel evolution happening in invasive ranges (Lucek *et al*., 2013; Latimer *et al*., 2019; McGoey *et al*., 2020). For this to occur, selection on *E. guttata* populations in Canterbury must be strong enough to overcome gene flow (Schmidt & Garroway, 2018). The selective gradient in Canterbury may be strong; the altitudinal gradient is similar in scale to some gradients in E. gutatta’s native range associated with ecotype formation (Hall *et al*., 2010; Popovic & Lowry, 2020), with significant differences in both temperature and precipitation across the gradient (Da Re *et al*., 2020; Williamson *et al*., 2023). Gene flow is also likely to be low in *E. guttata* populations due to its limited long-distance dispersal abilities (Waser *et al*., 1982). Relatively strong selection and weak gene flow may be driving differentiation even during this ongoing invasion process. The emerging variation may also become self-reinforcing, as increasing adaptive differentiation further limits gene flow (Lucek et al. 2013). Differentiation is also occurring in suites of traits, which is connected to strong evolutionary impacts (Lucek *et al*., 2013; Latimer *et al*., 2019; Felmy *et al*., 2022). In *E. guttata*’s native range altitudinal differentiation does not always happen in suites of traits due to the contrasting selection for drought resilience (DeMarche *et al*., 2020). As invasive plants *E. guttata* in Canterbury primary inhabit human-modified habitats where water is available year-round, adaptation may be more consistent in this range. Notably, similar differentiation has been seen in invasive *Erica lusitanica* in Canterbury (Mather & Williams, 1990), suggesting this may be a recurring pattern in plant invasions in the region.

### 4.2 Strong but Unspecialised Plastic Responses to Altitude

Canterbury *E. guttata* showed high levels of plasticity to growing environment in most traits, but there was little evidence that this plasticity was adaptive or that plastic responses varied between upland and lowland populations. Universally high plasticity is consistent with a general-purpose genotype invader (Matesanz *et al*., 2012) and with known high plasticity levels in *E. guttata* (Galloway, 1995; Murren & Dudash, 2012; Querns *et al*., 2022; Williamson *et al*., 2023). Consistently lower trait values were observed in the more abiotically challenging upland garden, which is indicative of passive rather than adaptive plasticity (Schneider, 2022). The specific adaptive significance of these high plasticity levels is therefore unclear from this study alone (Schneider, 2022), though earlier work in New Zealand *E. guttata* suggests high plasticity levels play a major role in facilitating invasive success (Williamson *et al*., 2023). There was little evidence of diverging plastic responses between origin groups. This mirrors earlier findings of chaotic variation in levels of plasticity among New Zealand *E. guttata* populations (Williamson *et al*., 2023). The two traits where we did find different levels of plasticity between origin groups may reflect selective differentiation based on altitude, with upland *E. guttata* populations having greater plasticity in the reproductive trait of flower number while lowland *E. guttata* populations had greater plasticity in the growth trait of height (Choler, 2001; van Kleunen & Fischer, 2008; Johansson *et al*., 2013). However, this is not conclusive, and different plastic responses were not seen across suites of traits which would indicate plasticity levels are evolving (Lucek *et al*., 2014; Felmy *et al*., 2022).

Traits which do not show any plastic change reveal important details about evolutionary success strategies, especially in highly plastic species (Matesanz *et al*., 2010). We identified a lack of plastic variation in all seed traits and in flower size, which is itself closely correlated with seed production (Kleunen & Ritland, 2004). These traits have a large genetic component in *E. guttata* (Robertson *et al*., 1994). Seed traits are closely related to success within a niche (Bufford & Hulme, 2021b). In *E. guttata*’s case, individual seeds have low survival odds (DeMarche *et al*. 2016), so seed traits may be genetically fixed to maximise seed number while maintaining consistently high germination rates (DeMarche *et al*., 2016; Rubin & Friedman, 2018). The lack of plastic variation between upland and lowland gardens suggests that these specific seed phenotypes maintain fitness across altitudinal environments (Murren & Dudash, 2012). Low plasticity was also seen in the different anthocyanin levels, which may reflect specific genetic colour morphs (Twyford *et al*., 2018). No plastic difference was seen in biomass, but this is likely a product of the autumn harvesting time.

### 4.3 The Importance of Adaptive Divergence Occurring Alongside High Plasticity

It is significant that this experiment identified adaptive differentiation to an altitudinal gradient happening in a general-purpose genotype invader alongside strong plastic responses to the same gradient. General-purpose genotypes often prevent genetic adaptation, even over much larger scales (Geng *et al*., 2007; Ross *et al*., 2009; Matesanz *et al*., 2020). This has been seen in *E. guttata* itself for thermal performance (Querns *et al*. 2022). Our experiment shows that general-purpose genotype invaders are not precluded from local adaptation. This is important because adaptation can directly increase invasiveness (Hock *et al*., 2019). High plasticity already facilitates global invasions of *E. guttata* (Querns *et al*., 2022); knowing genetic adaptation is still occurring during invasions of general-purposes genotype invaders reveals potential for future further increases in their invasive threat (Maron *et al*., 2007). Being able to mount adaptive genetic responses may also be important for future climate change responses (Richardson *et al*., 2014; Clements & Jones, 2021), as plastic responses alone may not be sufficient (Franks *et al*., 2014). Overall, this finding highlights more avenues for future survival and invasive success in already globally widespread general-purpose genotype invaders. Further research should focus on reciprocal transplants and genetic research to assess the strength of local adaptation in Canterbury *E. guttata*, identifying how common this pattern is in Canterbury plant invasions, and increasing our understand of how frequently local adaptation occurs in other general-purpose genotype invaders.

