## Supplementary Materials for "Adaptive Differentiation in the General-Purpose Genotype Invasive Plant *Erythranthe guttata*"

Table S1: Sampling locations of *Erythranthe guttata* across Canterbury

| Location | Origin Classification | Co-ordinates | Altitude (masl)* | Individuals in Lowland Garden | Individuals in Upland Garden |
|---|---|---|---|---|---|
| Boundary Stream | Upland | -44.053611, 170.133775 | 540 | 8 | 3 |
| Lake Pukaki | Upland | -43.973352, 170.130578 | 580 | 6 | 3 |
| Mt Cook Village | Upland | -43.738586, 170.097493 | 740 | 12 | 4 |
| Lake Pukaki (upper) | Upland | -43.977947, 170.129684 | 600 | 2 | 2 |
| Lake Ohau | Upland | -44.255570, 169.829824 | 540 | 9 | 4 |
| Lake Ohau | Upland | -44.227197, 169.816830 | 540 | 10 | 4 |
| Parsons Creek | Upland | -44.248745, 169.820028 | 540 | 9 | 4 |
| Lindis Pass | Upland | -44.517915, 169.727207 | 740 | 9 | 4 |
| Ahuriri River | Upland | -44.504069, 169.779595 | 560 | 8 | 4 |
| Ahuriri River – Bridge Campsite | Upland | -44.468110, 169.987688 | 420 | 9 | 4 |
| Arthurs Pass | Upland | -42.909925, 171.558565 | 900 | 4 | 2 |
| Goldney Saddle | Upland | -43.010212, 171.738836 | 560 | 5 | 2 |
| Lake Grasmere | Upland | -43.066453, 171.775265 | 580 | 6 | 2 |
| Cave Stream | Upland | -43.198465, 171.742201 | 640 | 4 | 2 |

| | | | | | |
|---|---|---|---|---|---|
| Castle Hill | Upland | -43.233576, 171.722958 | 760 | 3 | 2 |
| Lake Lyndon | Upland | -43.299449, 171.708934 | 840 | 6 | 2 |
| Windy Point | Upland | -42.583983, 172.384424 | 480 | 8 | 3 |
| Lewis Pass – Lawson Creek | Upland | -42.374901, 172.380720 | 880 | 6 | 1 |
| Speargrass Flat | Upland | -42.379065, 172.278443 | 540 | 4 | 2 |
| Mile Creek | Upland | -42.380595, 172.311326 | 560 | 2 | 2 |
| Marble Hill | Upland | -42.346827, 172.224481 | 460 | 5 | 2 |
| Lewis Pass | Upland | -42.378510, 172.399014 | 880 | 3 | 1 |
| Rolleston | Lowland | -43.609436, 172.384460 | 40 | 11 | 4 |
| Weedons Road | Lowland | -43.621170, 172.437448 | 20 | 10 | 4 |
| Lincoln | Lowland | -43.645329, 172.489597 | 10 | 7 | 4 |
| Bradley's Road | Lowland | -43.378991, 172.534542 | 40 | 6 | 3 |
| Jeff's Drain Road | Lowland | -43.393130, 172.597960 | 10 | 6 | 3 |
| Cust River | Lowland | -43.353259, 172.585718 | 20 | 10 | 3 |
| Silverstream | Lowland | -43.379538, 172.634206 | 10 | 6 | 3 |
| Kowai River – Mill Road | Lowland | -43.190815, 172.746087 | 10 | 5 | 3 |
| Amberly | Lowland | -43.159884, 172.727802 | 30 | 8 | 5 |
| Balcairn | Lowland | -43.193371, 172.675045 | 40 - 60 | 13 | 4 |

| | | | | | |
|---|---|---|---|---|---|
| Barbadoes Street | Lowland | -43.522931, 172.647076 | 5 | 11 | 4 |
| Avonside | Lowland | -43.504277, 172.682405 | 5 | 9 | 4 |
| Christchurch Botanical Gardens | Lowland | -43.527915, 172.627576 | 5 | 9 | 4 |
| Tinwald | Lowland | -43.915867, 171.726835 | 90 | 10 | 4 |
| Lake Hood | Lowland | -43.963325, 171.762389 | 60 | 10 | 4 |
| Methven Highway | Lowland | -43.857142, 171.731834 | 130 | 8 | 4 |

* Altitude determined from the 1:50,000 scale New Zealand Topographic Map, accessed from topomap.co.nz on 26/1/2024

*Supplementary Table S2:* Erythranthe guttata *phenotypic means and standard errors for origin groups at different garden locations, origin groups overall, and garden locations overall.*

| Trait | Upland origin *E. guttata* in lowland garden mean (SE) | Lowland origin *E. guttata* in lowland garden mean (SE) | Upland origin *E. guttata* in upland garden mean (SE) | Lowland origin *E. guttata* in upland garden mean (SE) | Upland Origin *E. guttata* overall mean (SE) | Lowland Origin *E. guttata* overall mean (SE) | *E. guttata* in lowland garden overall mean (SE) | *E. guttata* in upland garden overall mean (SE) |
|---|---|---|---|---|---|---|---|---|
| Bloom Period | 81 (1.53) | 74 (1.59) | | | 80 (1.40) | 74 (1.47) | 77 (1.11) | |
| Cold Damage | | | 4.1 (0.16) | 2.9 (0.17) | 4.1 (0.16) | 2.9 (0.17) | | |
| Corolla Width | 36.96 (0.470) | 37.16 (0.501) | 36.59 (0.596) | 37.27 (0.629) | 36.85 (0.444) | 37.2 (0.478) | 37.06 (0.344) | 36.93 (0.433) |
| Dry Weight | 50.126 (2.202) | 50.929 (2.373) | 50.543 (2.726) | 47.697 (2.885) | 50.252 (2.090) | 49.966 (2.274) | 50.529 (1.618) | 49.132 (1.984) |
| First Flower Date | 319 (1.82) | 334 (2.04) | | | 319 (1.82) | 334 (2.04) | 327 (1.37) | |
| Flower Number | 655.31 (25.90) | 525. 47 (27.07) | 286.43 (33.95) | 282.08 (35.28) | 550.50 (23.68) | 452.95 (25.20) | 595.14 (18.62) | 284.28 (24.51) |
| Height | 58.88 (1.92) | 72.76 (2.12) | 45.71 (2.25) | 47.97 (2.44) | 54.89 (1.84) | 65.37 (2.05) | 65.56 (1.43) | 46.83 (1.66) |
| Herkogamy | 2.8 (0.128) | 3 (0.134) | 3 (0.017) | 3.3 (0.172) | 2.82 (0.121) | 3.09 (0.128) | 2.87 (0.093) | 3.16 (0.119) |
| Leaf Anthocyanin Score | 1.5 (0.12) | 2.6 (0.14) | 1.7 (0.14) | 2.7 (0.15) | 1.6 (0.12) | 2.6 (0.13) | 2.1 (0.09) | 2.2 (0.10) |

| | | | | | | | | |
|---|---|---|---|---|---|---|---|---|
| Leaf Length | 55.25 (1.075) | 59.86 (1.131) | 33.69 (1.409) | 36.28 (1.463) | 48.72 (1.004) | 52.84 (1.066) | 57.56 (0.780) | 34.98 (1.016) |
| Leaf Width | 40.38 (0.830) | 47.27 (0.877) | 26.78 (1.093) | 31.07 (1.137) | 36.26 (0.760) | 42.44 (0.812) | 43.84 (0.604) | 28.9 (0.788) |
| Leaf Width-Length Ratio | 0.73 (0.0132) | 0.80 (0.0144) | 0.80 (0.0162) | 0.87 (0.0172) | 0.76 (0.0126) | 0.82 (0.0138) | 0.77 (0.0098) | 0.83 (0.0118) |
| Photosynthetic Rate | 7.96 (0.240) | 9.32 (0.249) | 8.65 (0.352) | 9.64 (0.369) | 8.17 (0.208) | 9.42 (0.218) | 8.41 (0.181) | 9.14 (0.282) |
| Seed Mass per Pod | 0.01667 (0.00054) | 0.01708 (0.00056) | 0.01524 (0.00073) | 0.01725 (0.00076) | 0.01624 (0.00051) | 0.01713 (0.00053) | 0.016876 (0.00039) | 0.01624 (0.00052) |
| Seed Number | 1219.6 (50.46) | 1284.8 (53.87) | 1138.2 (63.46) | 1262.9 (67.21) | 1195.0 (47.70) | 1278.3 (51.52) | 1252.4 (36.91) | 1200.0 (46.20) |
| Seed Size | 0.013866 (0.0004686) | 0.013795 (0.0005109) | 0.013906 (0.0005545) | 0.014377 (0.0005948) | 0.013878 (0.0004518) | 0.013968 (0.0004968) | 0.013830 (0.000347) | 0.014140 (0.0004067) |
| Stem Anthocyanin Score | 2.4 (0.19) | 3.7 (0.21) | 2.9 (0.21) | 4 (0.24) | 2.6 (0.19) | 3.8 (0.21) | 3.1 (0.14) | 3.5 (0.16) |

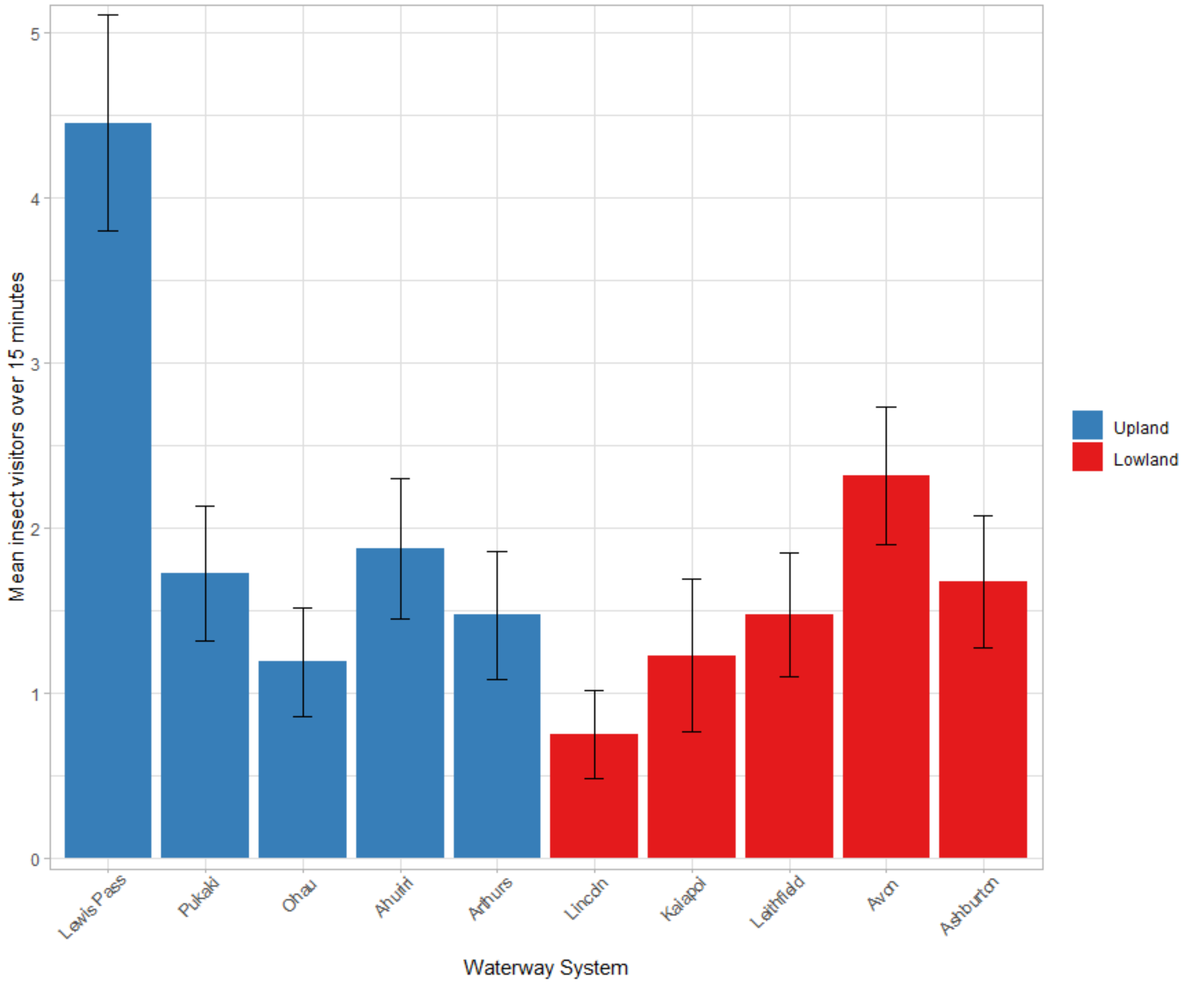


*Figure S1: Mean insect visits in 15 minute periods to* Erythranthe guttata *flowers in upland and lowland Canterbury populations, aggregated by waterway system from multiple sites per waterway. Lewis Pass was removed from statistical analyses as an outlier, as data could only be collected from a single atypical site.*
